# Predicting the global economic costs of biological invasions by tetrapods

**DOI:** 10.1101/2024.08.15.606318

**Authors:** Thomas W Bodey, Ross N. Cuthbert, Christophe Diagne, Clara Marino, Anna Turbelin, Elena Angulo, Jean Fantle-Lepczyk, Daniel Pincheira-Donoso, Franck Courchamp, Emma J Hudgins

## Abstract

Globalisation has steadily accelerated rates of biological invasions worldwide, leading to widespread environmental perturbations that often translate into rapidly expanding socioeconomic costs. Although such monetary costs can be estimated based on the observed effects of invasions, the pathways that lead invasive species to become economically impactful remain poorly understood. Here, we implement the first global-scale test of the hypothesis that adaptive traits that influence demographic resilience predict economic costs, using invasive terrestrial vertebrates as models given their rising impacts and well-catalogued characteristics. Our results reveal that total global costs of invasive tetrapods are conservatively in the tens of billions of dollars, with the vast majority due to damage costs from invasive mammals. These monetary impacts are predicted by longevity, female maturation age, diet and invasional pathway traits, although the directionality of predicted economic impacts also varied by trait across classes. Alarmingly, costs remain unknown for >90% of recorded established alien tetrapods worldwide, and across the majority of invaded countries. These huge socio-economic costs demonstrate the necessity of mitigating tetrapod invasions and filling knowledge gaps. Effective identification of traits predictive of costs among and within these groups can facilitate the prioritisation of resources to efficiently target the most damaging existing and emerging invasive tetrapod species.

## Introduction

Biological invasions — the establishment and spread of species outside their native ranges following human-mediated introductions (Blackburn et al. 2011) — are the cause of severe disruptions to ecological (Kumschick et al. 2014; Pyšek et al. 2020) and socioeconomic (Jones et al. 2017; Diagne et al. 2021; Bradshaw et al. 2024) systems globally, and a key driver of the global biodiversity decline (Brook et al. 2008, Doherty et al. 2016, IPBES 2023). Rapidly amplifying impacts of invasive alien species have been reported across a broad range of ecological networks, geographic regions, and socioeconomic systems (Bellard et al. 2016; Dick et al. 2017; Diagne et al. 2021). Therefore, syntheses of these impacts are crucial for rationalising policy design and motivating transboundary management actions (Dana et al. 2013; Lodge et al. 2016; Ahmed et al. 2023, IPBES 2023). They also achieve the critical goal of increasing societal knowledge and awareness — an essential but often missing requirement (Courchamp et al. 2017, IPBES 2023), given that rates of introduction and establishment of invasive alien species continue to rise, with no sign of abating (Seebens et al. 2017, 2021).

Across the globe, invasive tetrapods (i.e. amphibians, birds, mammals, and reptiles) are recognized as the cause of a range of concerning environmental, health and economic impacts (Clout & Russell 2008; Kraus 2015; Evans et al. 2020; Polaina et al. 2020; Wang et al. 2022; IPBES 2023).Their high establishment rates and broad range of impacts place these organisms among the most significant causes for conservation concern (Pysek & Richardson 2010; Bellard et al. 2016), particularly for their primary role in driving the extinction of endemic species and the erosion of functional diversity across islands worldwide (Blackburn et al. 2004; Drake et al. 2011; Doherty et al. 2016; McCreless et al. 2016; Bellard et al. 2017; Spatz et al. 2022). Cases where invasive species have triggered perturbations beyond recovery (i.e., tipping points) are abundant. For example, the elimination of much of Guam’s endemic avifauna by the brown tree snake (*Boiga irregularis*) (Rodda and Savidge, 2007). Likewise, the dominant role of invasive mammals and amphibians in the decline or extinction of multiple native species, and ongoing ecosystem destruction across Australia and New Zealand (Russell & Broome 2016; Stobo-Wilson et al. 2022; others *ad nauseum*); the global-scale loss of seabirds through the predatory impacts of invasive mammals (Jones et al. 2016). Invasive tetrapods also act as important reservoir hosts, and facilitate the vectoring of zoonotic pathogens and parasites (Crowl et al. 2008; Chalkowski et al. 2018), including the chytrid fungus implicated in the early stages of the current global. decline of amphibians (Schelle et al 2019).

In addition to their widely known environmental impacts, tetrapod invasions result in substantial financial expenditures, although these costs vary significantly across geographic regions, time periods and taxa (Diagne et al. 2021; Soto et al. 2022; Wang et al. 2022). For instance, rats (genus *Rattus*) cause substantial societal and health impacts to humans (Meerburg et al. 2009), resulting in significant control and eradication costs globally (Russell & Broome 2016; Bodey et al. 2022; Diagne et al. 2023). Likewise, feral pigs (*Sus scrofa*) produce huge agricultural, ecological and epidemiological costs globally (Bengsen et al. 2014, Risch et al. 2021). Among ectotherms, the damage caused by the brown tree snake to electrical and military infrastructure has incurred costs >$10 billion in the past 30 years (Soto et al. 2022); and coquí frogs (*Eleutherodactylus coqui*) reduce property values as a result of the noise pollution caused by their vocalisations (Kaiser & Burnett, 2014).

Despite the widespread and rapidly rising monetary costs exerted by invasive tetrapods, the pathway that introduced species traverse from introduction to economic impacts remains poorly understood. Therefore, developing a quantitative and integrative understanding of the dynamics underlying the past and present cost patterns is a critical step towards providing robust insights to predict — and then anticipate and mitigate — future such costs (Ahmed et al. 2022). In particular, given the role that life history traits play in the demographic resilience and rates of adaptability of populations, these fitness-relevant phenotypes have increasingly been recognized as important factors underlying invasion success (Sol et al. 2012; Capellini et al. 2015; Allen et al. 2017; Fournier et al. 2019; Street et al. 2023). Consequently, such traits are also anticipated to offer key predictors of economic costs, but remain to be widely harnessed (Evans et al. 2023). For example, ecological generalism is a consistent predictor of high impacts in taxa including birds (Sol et al. 2012; Evans et al. 2023; Marino & Bellard 2023) and reptiles and amphibians (Pili et al. 2024). Identification of traits conducive to high costs can therefore identify taxa and associated invasion pathways that warrant greatest current attention or preventative action (Turbelin et al. 2022; Cuthbert et al. 2023), thus enhancing management efficacy. Importantly, trait-based profiling is also used to underpin imputation approaches that quantify potential invasiveness, including the potential impacts of species that do not yet have an invasion history (Fournier et al. 2019; Pili et al. 2024). Despite this need to incorporate traits — and thus evolutionary and demographic insights — into the quantification of economic costs by invasive species, no such integrative studies exist.

Here, using the InvaCost database (Diagne et al. 2020), we implemented the first comprehensive global-scale study on the economic costs associated with invasive tetrapods. We then examined the utility of key ecological, life history and invasional traits of invasive tetrapod species for profiling the more costly invaders. Finally, we cross-referenced the reported socioeconomic costs with the most recent data on invasive tetrapod distributions (Seebens et al. 2020), highlighting how trait identification may facilitate or mitigate future cost increases and the extent of current knowledge gaps. Addressing these aims is an essential step towards developing predictive tools to inform much-needed policy and management actions.

## Materials and methods

### Cost data extraction and filtering

We extracted cost data from the latest version of the InvaCost database (version 4.1, open access at https://doi.org/10.6084/m9.figshare.12668570), a publicly available ‘living’ repository that compiles the monetary impacts of invasive species reported in English and 23 non-English language documents (Diagne et al. 2020; Angulo et al. 2021). InvaCost was created following a systematic, standardised methodology to collect information from peer-reviewed scientific articles, grey literature, stakeholders and expert elicitation. Following a thorough hierarchical screening of each source document for relevance, costs were extracted, standardised to a common currency (2017 US dollars (US$)) and associated with a range of descriptive fields pertaining to the original source (e.g. title, publication year), spatial and temporal coverage (e.g. period documented, study area), cost estimation methodology (e.g. reliability of values and acquisition method) and the cost estimates *per se* (e.g. nature and typology of cost). InvaCost thus represents the most up-to-date, comprehensive, and harmonised resource that records and describes economic cost estimates associated with biological invasions worldwide (Ahmed et al. 2023).

We applied a range of successive filtering steps (figure S1) using information from the descriptive columns within the InvaCost database (full details available at https://doi.org/10.6084/m9.figshare.12668570; see ‘Descriptors’ file). First, data pertaining to tetrapod taxa were obtained through selecting cost entries exclusively associated with one of the following classes: *Amphibia, Aves, Mammalia* or *Reptilia* (column ‘Class’). This resulted in a ‘full dataset’ containing 2375 cost entries representing a total of 107 species (7 amphibians, 20 birds, 61 mammals, 20 reptiles, Supplementary file 1: *full_dataset*). Second, to ensure the most robust and conservative synthesis of reported cost estimates, we subset this full dataset to include entries only if they were *(i)* successfully standardised from local currency to 2017 US$ (column ‘Cost estimate per year USD exchange rate’); *(ii)* of high (as opposed to low) reliability based on whether the approach used for cost estimation was traceable and reproducible (column ‘Method_reliability_refined’); and *(iii)* observed (as opposite to potential) costs, i.e. including incurred, but not projected, costs (column ‘Implementation’), while retaining estimates where empirically observed costs at small scales were then inferred over larger invaded areas or time periods. The resulting ‘robust dataset’ (Supplementary file 1: *robust_dataset*) contained 1764 cost entries. In order to homogenise the temporal occurrence of entries, all costs were standardised using the *expandYearlyCosts* function of the ‘invacost’ R package (Leroy et al. 2022) to provide annual cost estimates for all entries based upon the start and end years provided in the database (columns ‘Probable starting year adjusted’ and ‘Probable ending year adjusted’). A single cost entry without a starting year was then omitted. The resulting ‘expanded dataset’ (Supplementary file 1: *expanded_dataset*) contained 3185 rows of costs representing a total of 92 species (7 amphibians, 15 birds, 53 mammals, 17 reptiles). Finally, for consideration of predictive traits, only cost estimates attributable to individual species were retained. This excludes, for example, costs that could not be accurately partitioned among multi-species control or eradication actions from the information available. This latter analysis was thus conducted on 2478 rows of costs across 80 tetrapod species (Supplementary file 1: *-expanded_trait_dataset*).

### Cost estimations and distributions

Total cost estimates were obtained by summing values in the ‘Cost estimate per year 2017 exchange rate’ column from the expanded dataset. We examined the magnitude of these costs over several categories pertaining to different descriptive fields (for full details see Table S1), namely: *type of cost* (Damage, Management or Mixed), *spatial distribution* of costs across continental regions, *taxonomic distribution* of costs across Classes and Genera and *impacted sectors* (Authorities-Stakeholders, Agriculture, Environment, Fisheries, Forestry, Health, Public and social welfare, or mixtures of these) (table S1, Diagne et al. 2020).

### Association between traits and recorded costs

In order to examine the potential effect of species traits and invasion-related metrics on the amount and total costs, we collated species-level information from multiple databases including: main diet, level of habitat generalism, body mass, female maturation, lifespan, time to independence, reproduction, date of first invasional record, invasion pathway (table 1 for further details).

**Table 1:** Ecological, life history and invasional variables included in phylogenetically-controlled analyses of predictive traits for economic costs.

| Trait | Trait Type | Variable Type | Description |
| --- | --- | --- | --- |
| Diet<sup>1,2,3</sup> | Ecological | Categorical | Main diet category (herbivore, omnivore, carnivore) based on the dominant dietary type (>50%) consumed (*cf* Marino et al. 2022) |
| Habitat Generalism<sup>3,4,5,6</sup> | Ecological | Binary | Specialist (occupies a single or very similar habitat types) or Generalist (occupies multiple differing habitat types) |
| Body Mass<sup>3,7</sup> | Life History | Continuous | Mean adult body mass (grams) |
| Female Maturation<sup>3,7</sup> | Life History | Continuous | Mean time to female reproductive maturity (months) |
| Lifespan<sup>3,7</sup> | Life History | Continuous | Mean lifespan (years) |
| Time to Independence<sup>3,7</sup> | Life History | Continuous | Weaning or fledging age (years) |
| Reproduction<sup>3,7</sup> | Life History | Continuous | A composite of the number of offspring/clutch size per reproductive event and the number of reproductive events/year |

| First Record<sup>8</sup> | Invasional | Continuous | Date of first record in each invaded country with economic costs reported |
| --- | --- | --- | --- |
| Invasion Pathway<sup>9</sup> | Invasional | Categorical | Intentional (e.g. release, escape), Unintentional (e.g. stowaway, corridor disperser) or Unknown |

<sup>1</sup>Pigot et al. (2020), <sup>2</sup>Wilman et al. (2014), <sup>3</sup>Pincheira-Donoso 2017, <sup>4</sup>Tobias et al. (2022) Ecol Lett, <sup>5</sup>Faurby et al. (2018) Ecol, <sup>6</sup>Uetz et al. (2022), <sup>7</sup>Mhyrvold et al. (2015) Ecology, <sup>8</sup>Seebens et al. (2020), <sup>9</sup>Turbelin et al. (2022) Biol Inv

Central to robust multi-species comparative analysis is the need to control for phylogenetic non-independence as a result of relatedness through shared ancestry (Stone et al. 2011). For each taxon, we obtained phylogenetic information using the phylogenetic eigenvectors computed from 1,000 trees based on phylogenies from www.vertlife.org (Jetz et al. 2012; Tonini et al. 2016; Jetz & Pyron 2018; Upham et al. 2019). Eigenvectors extracted from principal component analyses represent the variation in the phylogenetic distances among species, with the first 10 eigenvectors explaining more than 50% of this phylogenetic variance. We thus used the phylogenetic eigenvectors as a control in all trait models, fitting separate models for each tree, and thus set of eigenvectors, resulting in 1000 models per taxonomic class. These terms can also be interpreted as the effect of relatedness on economic impact, independent of the traits included in the model. We included up to three eigenvectors across our models to balance the limited degrees of freedom in our relatively small sample sizes (and therefore, limited number of additional degrees of freedom) with an appropriate amount of phylogenetic variation captured in our models. We removed any eigenvectors that were highly correlated within our sample.

The decade in which a cost was incurred was added as an additional predictor in explanatory models to control for consistent changes in cost magnitude over time across records (for instance, due to growing awareness of invasive species’ impacts (Seebens et al. 2017; Diagne et al. 2021). To allow for non-linear changes in the cost magnitude over time, we implemented generalised additive models (GAMs) using the *mgcv* package (Wood 2011), with decade as a smoother term. All other variables were incorporated as linear terms.

Models were run with total costs as the independent variable and the traits outlined above as explanatory variables. Parameter estimates and mean p-values were derived from the mean and 95% quantiles of 1000 model runs. Each taxonomic class was run separately, with an additional model using data from all tetrapods combined.

### Taxonomic and cost completeness

We used the most up-to-date distributional information for all known invasive species globally (sTwist, Seebens et al. 2020) to assess the completeness of the expanded, filtered dataset relative to the distributions of invasive tetrapods. We extracted the total set of known invasive tetrapods from sTwist, together with their date(s) of introduction and established invaded ranges. We also cross-referenced our InvaCost dataset with the Global Register of Introduced and Invasive Species (GRIIS, www.griis.org). GRIIS is a database of invasive species’ temporal and spatial impacts maintained by the Invasive Species Specialist Group of the IUCN. It records expert evidence of any form of ecological impacts from invasive species across locales. Here we specifically compared the records of species within our economic cost dataset with those where impact (a binary yes/no within GRIIS) had been identified. We used the *‘*countrycode’ R package to convert country records to commensurate codes in order to merge the databases (Arel-Bundock et al. 2018). Note that this package includes some overseas territories and other locales as separate units, although they may not qualify as countries under stricter definitions. We considered a species missing from InvaCost relative to sTwist or GRIIS if InvaCost did not report a cost for any country listed as part of the invader’s range within the other database. We thus determined the extent of missing established species from InvaCost (i.e. known invaders with unknown costs) among invasive tetrapods.

All analyses were conducted in R version 4.1.0.

## Results

### Global patterns of monetary costs

The total reported economic costs of invasive tetrapods was $55.2 billion from 3,185 expanded cost entries. The vast majority (∼85%) of this global cost was incurred through resource damages and losses ($47.0 billion), with the highest geographic totals incurred in North America ($28.0 billion) and Oceania ($11.1 billion) (figure 1). Of the $3.1 billion (∼6%) management spend globally, $3.0 billion was spent post-invasion, with two orders of magnitude less invested pre-invasion ($52.6 million). Mammals dominated these costs ($52.5 billion), with birds ($1.4 billion), reptiles ($1.2 billion) and amphibians ($0.05 billion) all accounting for far lower values (figure 1). The costliest genera were also most often mammals, with *Rattus* ($30.8 billion), *Felis* ($6.2 billion), *Sus* ($4.3 billion), *Canis* ($2.4 billion), *Oryctolagus* ($1.9 billion) and *Boiga* ($1.2 billion) all contributing over $1 billion each, and with all other invasive tetrapod genera combining to ∼$2.8 billion in total. Among activity sectors, costs attributable to specific sectors revealed agriculture to be impacted to the greatest extent ($8.2 billion), followed by authorities-stakeholders ($6.0 billion), health ($4.7 billion), the environment ($2.6 billion) and public and social welfare ($1.4 billion); with remaining specific sectors each incurring under $1 billion in costs (figure 1). Notably, costs spread indistinguishably across authorities-stakeholders and the environment totalled $23.8 billion.

**Figure 1.**
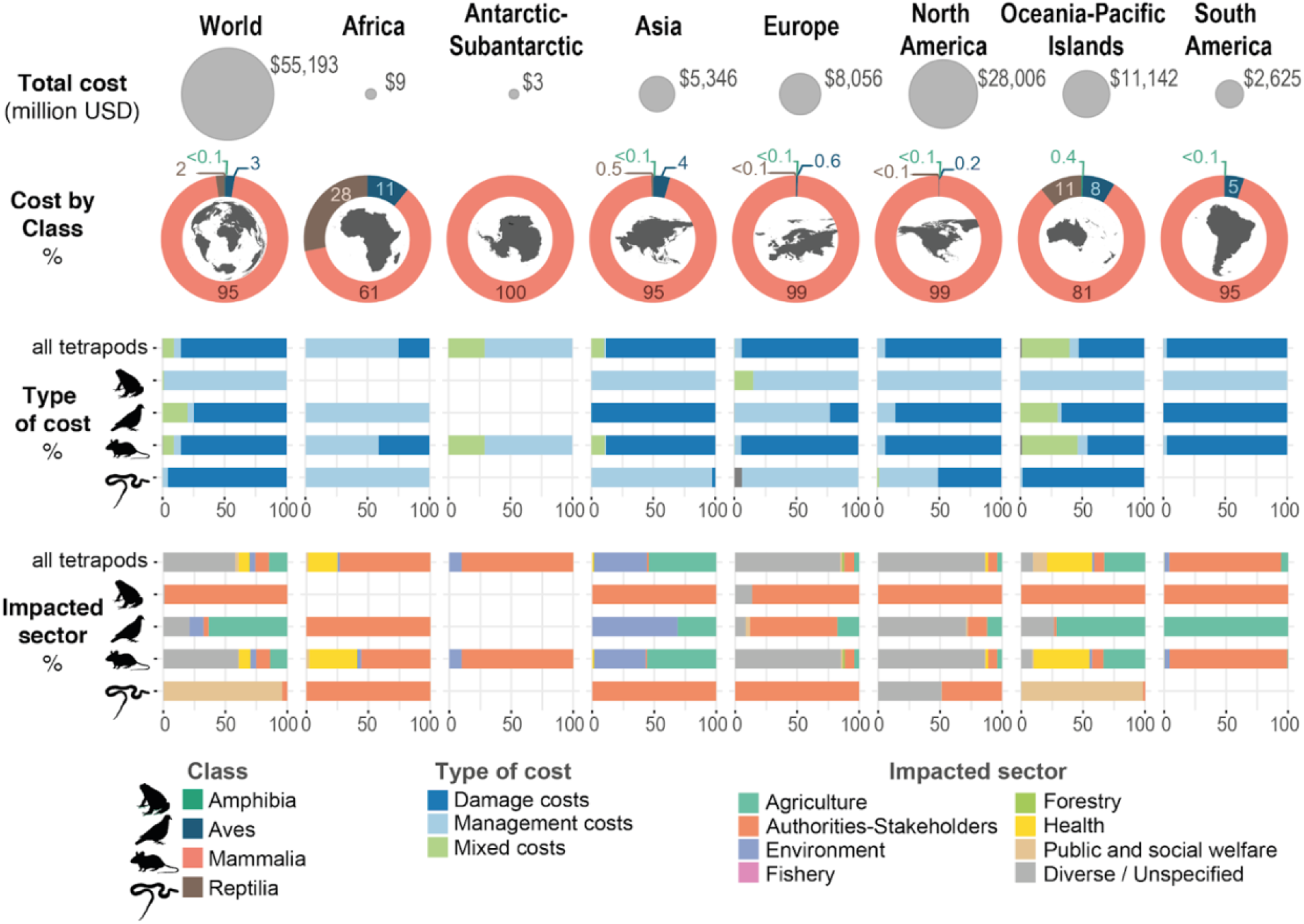
Distributions of economic costs of invasive tetrapods according to geographic regions, taxonomic classes, cost types and impacted sectors.

### Predictive Traits

When examined across all invasive tetrapods together (figure 2), higher economic costs were significantly positively associated with taxa possessing higher maximum longevity and those introduced via multiple and unknown pathways; while costs were significantly negatively associated with the age of female maturity. However, the key traits that predicted economic costs differed substantially between tetrapod taxonomic classes (figure 2). Amphibians had no significant non-phylogenetic predictors of costs. For birds, there was a limited set of significant predictive traits, with greater economic costs associated with greater longevity and earlier time to independence. For Mammals, higher economic costs were significantly associated with carnivorous and herbivorous diets, higher fecundity, and being introduced via intentional means, with no traits significantly predictive of lower costs. Carnivorous reptiles and those with later female maturity had increased costs, and phylogenetic signal was strong across both reptiles and amphibians (figure 2).

**Figure 2:**
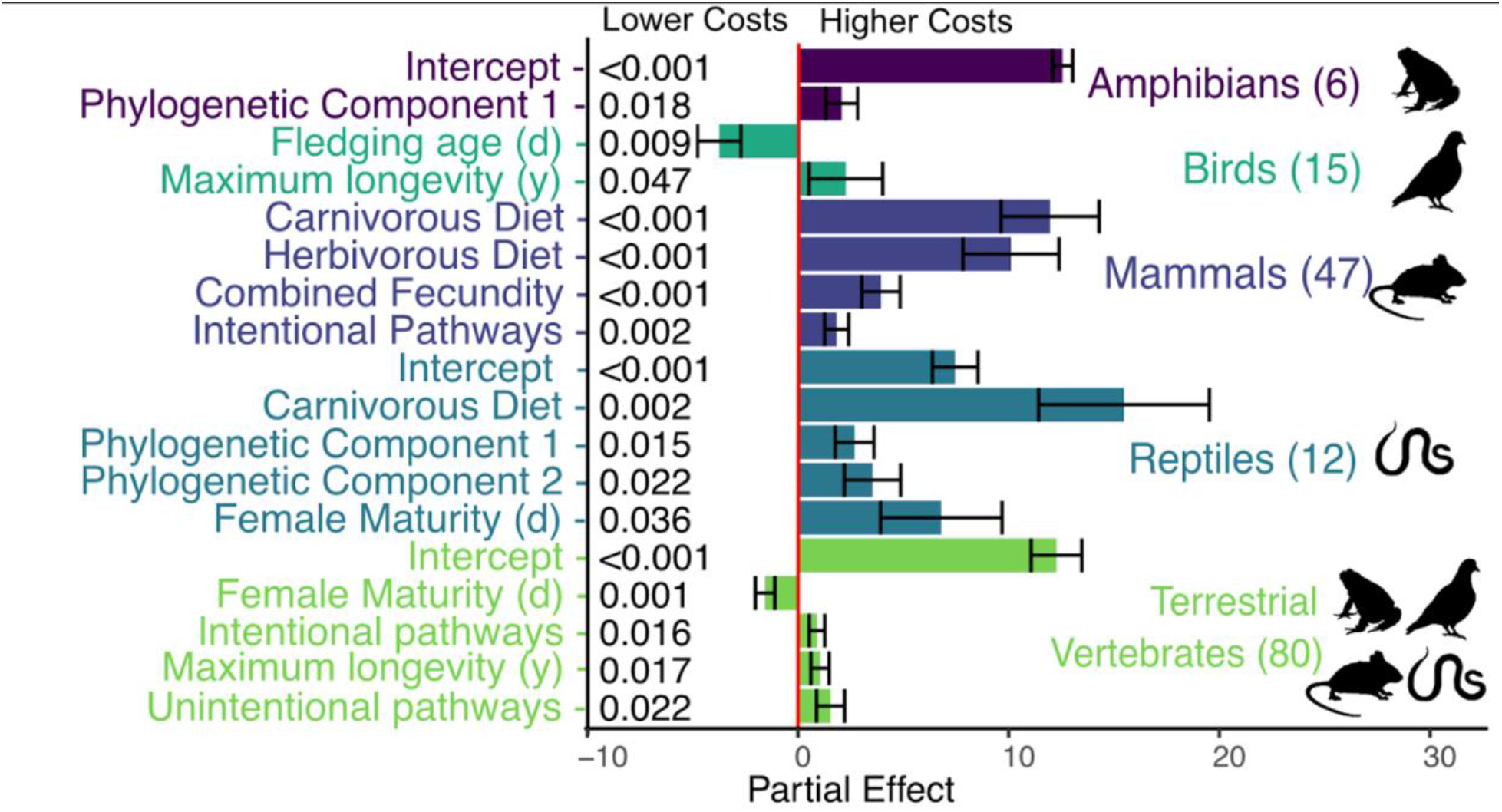
Partial effects plot showing significant trait predictors of costs on a logarithmic scale, with p-values shown to the left of each bar and results averaged over 1000 sampled phylogenetic eigenvectors. Partial effects of dietary components (Herbivorous and Carnivorous) are measured relative to Omnivorous in the case of Reptiles and Unknown in the case of Mammals. Pathway components (Intentional, Multiple, and Unintentional) are measured relative to Unknown pathways. The decadal smoother term was significant with a mean empirical degrees of freedom (edf) of 1.81 and a mean p-value of 0.029 for the all tetrapods model. For amphibians: smoother edf <0.001, p = 0.85; birds: smoother edf = 1.04 p = 0.0054; mammals: edf = 2.11, p = 0.0011; reptiles: edf <0.0001, p=0.49. The number of unique species in each group is shown in parentheses.

### Database Alignment

There were substantial differences among the records of invasive tetrapod species within distributional (sTWIST), impactful (GRIIS) and economic (InvaCost) databases (figure 3, table S2). Principally, these demonstrate a wide array of invasive species which, although recorded as present in many countries (sTWIST) or with ecological impact (GRIIS), still lack records of economic impacts within the same locales (InvaCost). Interestingly, only just over half of all species with reported economic costs are present in sTwist. InvaCost also contains evidence of socioeconomic impacts for a substantial number of invasive tetrapods that do not have ecological impacts recorded within GRIIS.

**Figure 3:**
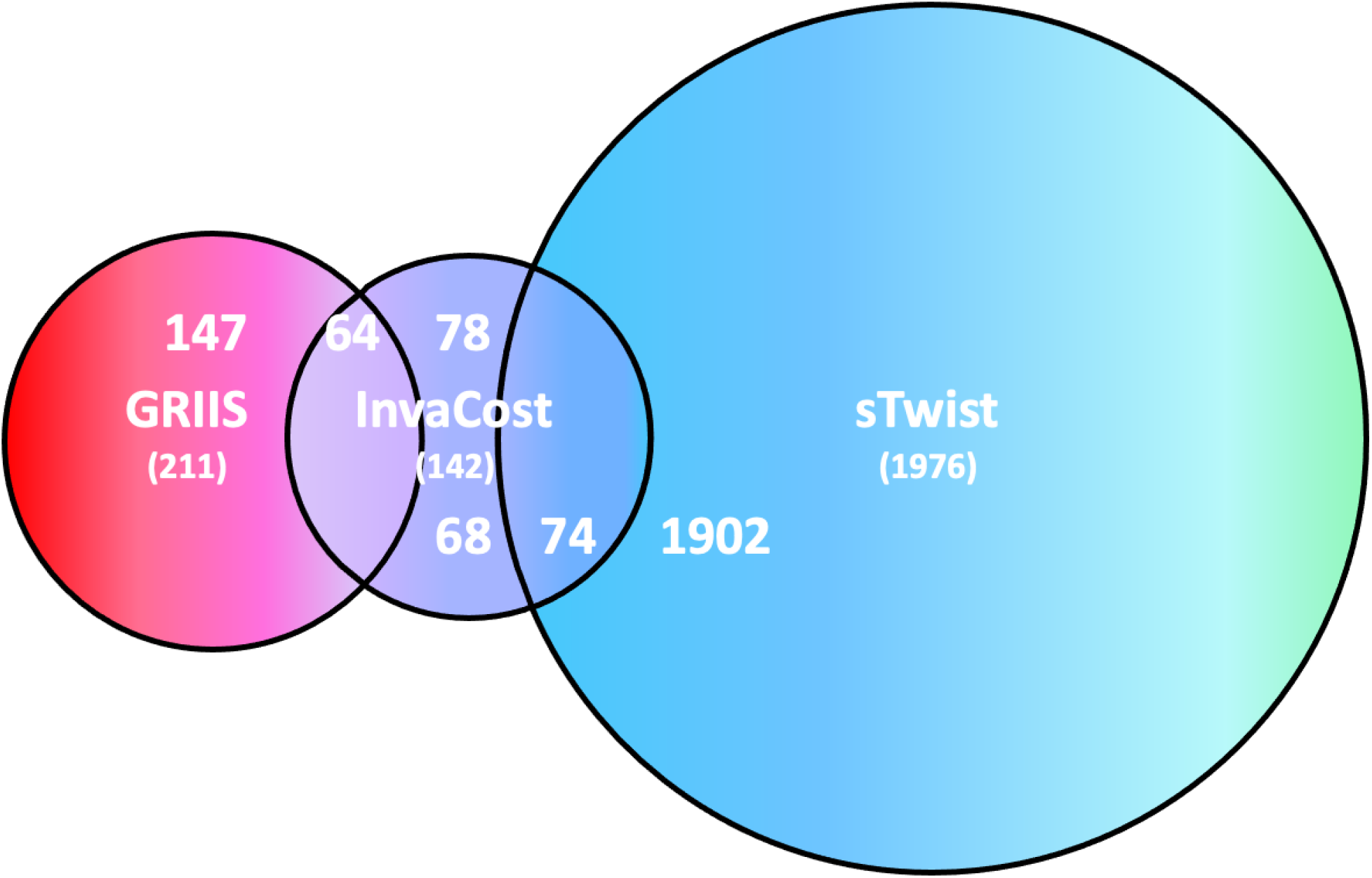
Comparative diagram (scaled by total record size) of the overlap in species records between databases on the economic cost of invasive tetrapods (InvaCost), records of their ecological impact (GRIIS) and their distribution (sTwist). InvaCost contains records directly attributable to 142 species, 64 of which are in common with the 211-species rich GRIIS dataset (leaving 78 only present in InvaCost, i.e. with identified socioeconomic but not ecological impacts), and 74 of which are shared with the 1976-species rich sTwist dataset.

## Discussion

Our study provides the first integrative comparative analysis of the global economic costs, and of the traits that predict these impacts, of invasive species from across the tetrapod tree of life. Our findings reveal substantial regional and taxonomic differences in reported economic costs, skewed to a few extremely costly genera, geographic regions, and sectors. This is reflected in the clear disparities in records of species across databases, with over 90% of known invasive alien tetrapods having no reported economic costs. However, our analyses were able to identify a set of key traits — notably maximum longevity, age of female maturation and principal diet — as significant predictors for economic costs across multiple tetrapod classes, although with variation in directionality.

### Global patterns of monetary costs

The recorded global costs of invasive tetrapods are huge (Diagne et al. 2021; Soto et al. 2022; Wang et al. 2022), second only to invasive alien insects worldwide (see ‘living figure’, Diagne et al. 2021). However, these costs vary immensely across tetrapod classes (figure 1). The overwhelming majority of costs were attributable to mammals, with some hyper-costly genera and species (*cf* Heringer et al. 2021) including rats, feral cats (*Felis catus*) and pigs accounting for much of the total, and only one non-mammalian hyper-costly species (the brown tree snake). Mammal invasion economic impacts have been shown to be significantly overrepresented compared to global non-native species richness, further indicating a high impact propensity identified from that class (Cuthbert et al. 2024).

Geographic disparities were also clear, with a data bias towards regions with greater available research funding, i.e. North America, Europe and Oceania (predominantly New Zealand and Australia) (Bellard & Jeschke, 2016; Diagne et al. 2021). Globally, damage costs far outweighed those of management, with many costs attributed to agricultural damage incurred chiefly by rodents (figure 1). Management efforts were significantly skewed towards post-invasion reactive responses such as eradications (Oppel et al. 2011; Veitch et al. 2019), with two orders of magnitude less invested in preventative, and relatively more efficient, pre-invasion actions that can reduce overall costs of inaction (Ahmed et al. 2022; Cuthbert et al. 2022). Interestingly, while small in comparison to the overall total, costs for amphibians were almost exclusively management-focussed, and the great majority of costs for herpetofauna were borne by authorities, stakeholders and other public sectors, in a clear contrast with the other tetrapod orders. Management of biological invasions has been found to be among the most effective conservation tools for bending the curve of biodiversity loss and ecosystem impacts (Ahmed et al. 2022; Langhammer et al. 2024), and so we urge improved investments to mitigate invasive species’ impacts across taxa.

### Predictive traits

We found that variables from all three trait types (ecological, life-history, and invasional) were important predictors across multiple taxonomic permutations (figure 2). Invasional pathways can shed light on the probable extent of propagule pressure, with regular deliberate or accidental transport of individuals increasing the likelihood of establishment (Hulme et al. 2008; Capellini et al. 2015; Early et al. 2016; Toomes et al. 2022). We found increased costs associated with intentional transport of mammals specifically. Deliberate introductions of various ungulates and lagomorphs as food or hunting resources (Lever et al. 1985; Soto et al. 2024a) and predatory mammals as intended biological controls (King 2017), have produced a wide range of global economic costs alongside ecological impacts (Diagne et al. 2021; Wang et al. 2022; Soto et al. 2024b). Indeed, intentionally introduced taxa are often overrepresented in global biological invasions (Briski et al. 2024). However, for tetrapods as a whole, both intentional and unintentional pathways of introduction were associated with higher economic costs than those with unknown pathways. This likely reflects the mechanisms outlined above, alongside other movements including the pet trade and redistribution of commensal species (Clout & Russell 2008; Soto et al. 2022; Toomes et al. 2023). Importantly, lower costs could be reflective of a subset of species subject to less research effort due to any combination of their invasion history, perceived impact or invaded locations, rather than a guarantee of these species’ lower economic impacts. Specific pathways have not previously been directly connected with the severity of ecological impacts in terrestrial vertebrates, albeit domestication and the pet trade have been identified as markers for impact risk (Gippet & Bertelsmeier 2021; Wang et al. 2024), with taxa kept or adapted to close association with humans more likely to be introduced and to have substantial ecological and economic effects (Jeschke & Strayer 2006; Soto et al. 2024a).

As well as species’ invasional history, we considered multiple ecological and life history traits. Having controlled for phylogeny, we found that maximum longevity and measures associated with fecundity and female maturation time were significant predictors of costs across multiple classes. Population dynamics are reflective of the filters imposed during the progress of invasions (Blackburn et al. 2011; Leung et al. 2012; Ahmed et al. 2022; Chapple et al. 2022) but, importantly, economic impact is independent of organism invasiveness as these are separate components of the invasion process, and economic costs can occur at any stage during an invasion (Marino & Bellard 2023; Soto et al. 2024b). Faster life histories with rapid maturation have repeatedly been found not to be predictive of more successful establishment probability, and hence ecological impact, in invasive mammals and birds (Sol et al. 2008, 2012; Capellini et al. 2015) — contrasting with the finding here with more rapid fledging time being predictive of greater economic costs in birds. Higher fecundity, as reflected in greater annual reproductive output in mammals and later female maturation time in reptiles (given the capacity for larger females to produce more young) was associated with increased costs, highlighting potentially strong links between these aspects of population dynamics and economic impacts. Diets also had class specific predictive capacity, again suggesting the potential for important predictive differences between endo- and ectothermic species. In contrast, when considering ecological impacts, diet was found to be an important predictor only for birds (Evans et al 2023; Wang et al 2024), again highlighting a potential disconnect between economic and ecological impact assessments. Overall, these results confirm that the directionality of predictive traits can be class-dependent, emphasising the benefit of a robust understanding of related species’ ecology and demography (Ricciardi et al. 2013, 2021).

There is an increasing focus on the use of traits to aid with a range of management approaches from species profiling to restoration (Laughlin 2014; Capellini et al. 2015; Fournier et al. 2019; Ricciardi et al. 2021; Latombe et al. 2022; Streit & Bellwood 2022; Cuthbert et al. 2023; Pili et al. 2024). Our ability to determine key predictive traits for economic costs are derived from living databases, thus increasing insight and predictive power will be gained as further information is accrued. For example, knowledge gaps surrounding invasive amphibians and reptiles (Soto et al. 2022, but see Pili et al. 2024) and economic impacts prior to noticeable damage costs (Diagne et al. 2021; Cuthbert et al. 2022) require novel research and investigation for more species in more places. However, this variation in available data is a common problem in invasion science, whether through a lack of detailed study, investigatory or publication biases, or invasional lags, and it is important to make assessments based on the available information, particularly where relationships are complex but actions and decisions are necessary and can have significant financial and ecological implications (Leung et al. 2012; Essl et al. 2020; Christie et al. 2020; Nunez et al. 2021; IPBES 2023).

### Database alignment

While databases compiling the distribution, ecological and economic impacts of invasive species would not be expected to completely align due to potentially multi-decadal publication or invasional timelags (Duncan 2021; Leroy et al. 2022), there was a much greater degree of dissimilarity than we anticipated. While a small number of species and costs may have been omitted here due to a lack of species-specific resolution for multi-species management actions or minor differences in classifications between external territories and governing countries, there remains a clear absence of information on the economic impacts of many invasive tetrapods. Comparison to the sTWIST database (the most complete record of global alien incursions) showed that 1902 known established alien tetrapods and approximately 200 invaded locations lacked reported costs. This clearly demonstrates the extent of socioeconomic cost underestimation (Henry et al. 2023; IPBES 2023), with even species with reported costs still lacking values from multiple locations within invaded ranges.

Similarly, the majority (70%) of invasive tetrapod species with known ecological impacts in GRIIS had no economic impacts recorded in InvaCost. In tandem though, many species with economic impacts were not captured in GRIIS, highlighting potentially strong asynchrony between ecological and economic impact research or the lack of attribution of economic impacts to invasive status (Vaissière et al. 2022). While multiple methods exist for the quantification of ecological impacts in economic terms (e.g., revealed/stated preference methods; Hanley and Roberts, 2019), such assessments are lacking for most tetrapods (Evans et al. 2023). These disparities also raise questions about the extent to which data and risk analyses are identified within, and shared across, international borders or among regional agencies (Turbelin et al. 2022; Bodey et al. 2023).

Disparities also likely result from a number of societal and terminological discrepancies around invasive tetrapods. These include those with both costs and benefits (e.g., introduced quarry animals) that produce multiple viewpoints on the relative extent of impacts (Shackleton et al. 2019; Kourantidou et al. 2022; Carneiro et al. 2024). Similarly, species with long invasion histories are often considered part of their ecological communities, and may be considered ‘pests’ rather than invasive, and thus conflated with morphologically similar native ‘pests’ (Diagne et al. 2023; Soto et al. 2024a). Discrepancies in cost attribution can also occur across research fields, with values directly associated with an invasive species in an ecological setting, but subsumed within medical or welfare outcomes within a health context. This is important as this often concerns hypercostly species (for example, invasive rodents with cross-sector impacts on native species, agriculture and public health [Witmer & Shiels 2018; Diagne et al. 2023]), so interdisciplinary approaches are essential to ensure costs for such species are fully recognised and effectively collated.

## Conclusions

We provide the first evidence that species’ traits can be harnessed to forecast economic impacts. In the future, these relationships could be applied to extrapolate and impute costs and impacts for missing invasive species and locations based on those with similar trait profiles, or to profile potential novel economically costly invaders (Fournier et al. 2019; Latombe et al. 2022; Henry et al. 2023; Pili et al. 2024). Such efforts could include incorporating the rapidly expanding global impacts of biological invasions into the planetary boundaries framework (Steffen et al. 2015), a key method for identifying catastrophic anthropogenic threat thresholds from which they are currently overlooked. All such efforts will be enhanced by further cost reporting and modelling that contributes to a more comprehensive picture of the true costs of invasive tetrapods (and other taxonomic groups), bolstering predictive capacities and informing and streamlining management strategies. Such prioritisation exercises are valuable for guiding decision making concerning both risk assessment and applied adaptive management given the inevitable presence of financial and logistical constraints (Carrasco et al. 2010; Leung et al. 2012; Booy et al. 2020; Carter et al. 2022). They are also essential for revealing the true level of threat provided by this global anthropogenic impact.

## Acknowledgements

The authors acknowledge the French National Research Agency (ANR-14-CE02-0021) and the BNP-Paribas Foundation Climate Initiative for funding the Invacost project that allowed the construction of the InvaCost database. The present work was conducted following a workshop funded by the AXA Research Fund Chair of Invasion Biology and is part of the AlienScenario project funded by BiodivERsA and Belmont-Forum call 2018 on biodiversity scenarios. RNC acknowledges funding from the Leverhulme Trust (ECF-2021-001). CD was funded by the BiodivERsA-Belmont Forum Project “Alien Scenarios” (BMBF/PT DLR 01LC1807C). JFL would like to thank the Auburn University College of Forestry and Wildlife Sciences for travel support to attend the Invacost workshop.

## CRediT authorship contribution statement

TWB: Conceptualisation, Data Curation, Formal analysis, Visualisation, Writing – original draft, Writing – review & editing

EJH: Data Curation, Formal analysis, Visualisation, Writing – review & editing

CD: Conceptualisation, Data Curation, Writing – review & editing

RNC: Formal analysis, Visualisation, Writing – review & editing

CM: Data Curation, Writing – review & editing

EA: Data Curation, Writing – review & editing

JF-L: Data Curation, Writing – review & editing

AT: Data Curation, Visualisation, Writing – review & editing

DP-D: Data Curation, Writing – review & editing

FC: Conceptualisation, Funding Acquisition, Writing – review & editing

## Data Availability Statement

The original data considered are openly accessible at doi.org/10.6084/m9.figshare.12668570. The final dataset used for our analyses are available as a supplementary material file.

---

**Figure S1.**
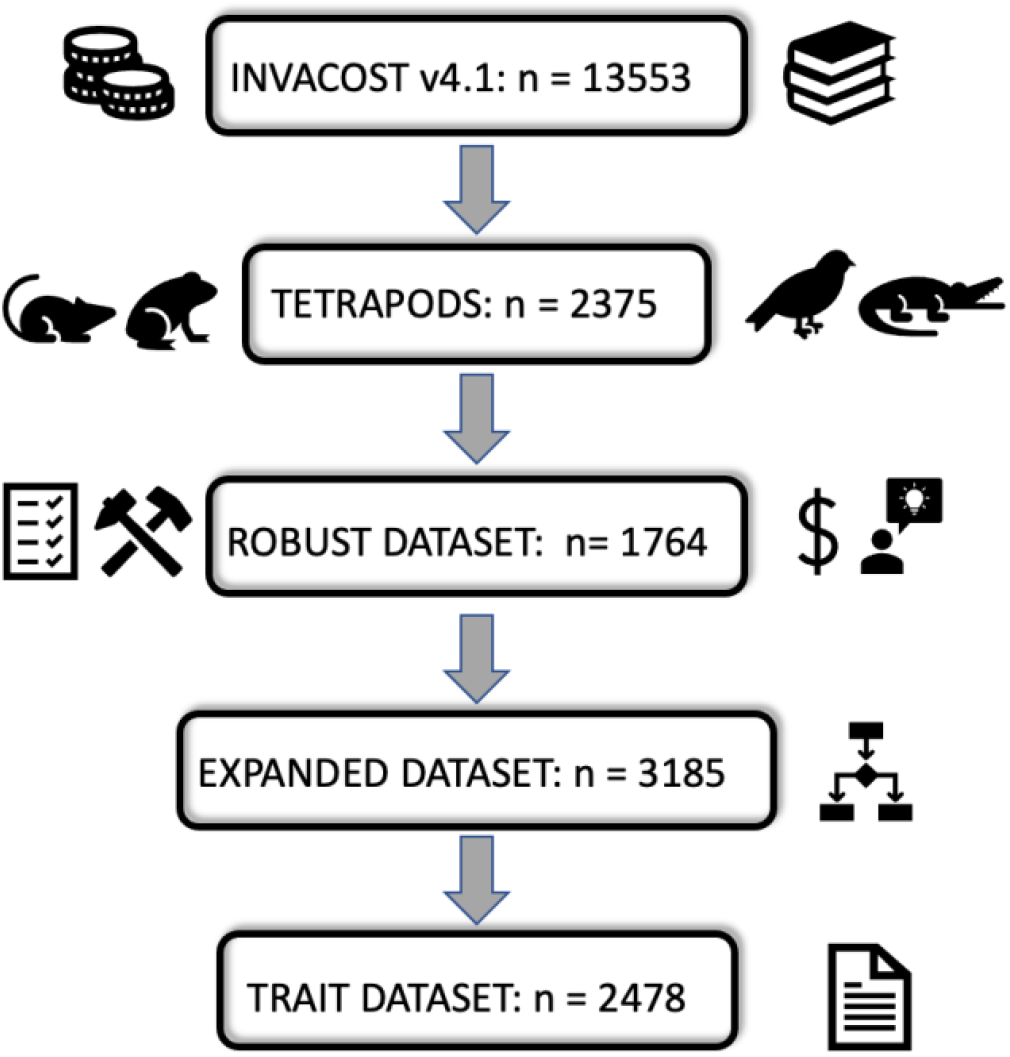
Filtering steps applied to the full InvaCost database to produce datasets for analyses. Sample sizes are reported in terms of rows of data retained from the full database and then following expansion to yearly quantities. Note cost distributions are assessed from the Expanded Dataset, whereas trait-cost associations are necessarily only analysed from the Trait Dataset where all costs could be attributed to specific tetrapod species.

**Table S1:** Categorisation of costs across different descriptive parameters from the InvaCost database.

| Distribution Category | Descriptive Column Used | Divisions |
| --- | --- | --- |
| Cost Type | Type of cost merged | *Damage*: losses due to direct or indirect impacts such as yield losses, repairs, or income reduction<br>*Management*: resources allocated to avoiding or reducing impacts such as control, research or mitigation policies<br>Management costs were further subdivided as:<br>*Reactive*: post-invasion responses e.g. control, eradication<br>*Proactive*: pre-invasion actions e.g. investment in prevention or biosecurity<br>*Knowledge funding*: investment in research and communications<br>*Mixed*: costs that simultaneously include both damage and management components or that cannot be attributed to a single cost division |
| Spatial Distribution | Geographic region | Africa, Asia, Europe, North America [including Central America], South America, Oceania [including Pacific Islands] |
| Taxonomic Distribution | Class and Genus | - |
| Impacted Sector | Impacted_sector | Authorities-Stakeholders, Agriculture, Environment, Fisheries, Forestry, Health, Public and social welfare, or Mixed (inseparable combinations of sectors) |

## References

Ahmed DA et al. 2022. Managing biological invasions: the cost of inaction. Biological Invasions 24:1927–1946.

Ahmed DA et al. 2023. Recent advances in availability and synthesis of the economic costs of biological invasions. BioScience 73:560–574.

Allen WL, Street SE, Capellini I. 2017. Fast life history traits promote invasion success in amphibians and reptiles. Ecology Letters 20:222–230.

Angulo E et al. 2021. Non-English languages enrich scientific knowledge: The example of economic costs of biological invasions. Science of The Total Environment 775:144441.

Arel-Bundock V, Enevoldsen N, Yetman C. 2018. countrycode: An R package to convert country names and country codes. Journal of Open Source Software 3:848.

Bellard C, Cassey P, Blackburn TM. 2016. Alien species as a driver of recent extinctions. Biology Letters 12:20150623.

Bellard C, Jeschke J m. 2016. A spatial mismatch between invader impacts and research publications. Conservation Biology 30:230–232.

Bellard C, Rysman J-F, Leroy B, Claud C, Mace GM. 2017. A global picture of biological invasion threat on islands. Nature Ecology & Evolution 1:1862–1869.

Bengsen AJ, Gentle MN, Mitchell JL, Pearson HE, Saunders GR. 2014. Impacts and management of wild pigs S us scrofa in A ustralia. Mammal Review 44:135–147.

Blackburn TM, Cassey P, Duncan RP, Evans KL, Gaston KJ. 2004. Avian Extinction and Mammalian Introductions on Oceanic Islands. Science 305:1955–1958.

Blackburn TM, Pyšek P, Bacher S, Carlton JT, Duncan RP, Jarošík V, Wilson JRU, Richardson DM. 2011. A proposed unified framework for biological invasions. Trends in Ecology & Evolution 26:333–339.

Bodey TW, Angulo E, Bang A, Bellard C, Fantle-Lepczyk J, Lenzner B, Turbelin A, Watari Y, Courchamp F. 2023. Economic costs of protecting islands from invasive alien species. Conservation Biology 37:e14034.

Booy O et al. 2020. Using structured eradication feasibility assessment to prioritize the management of new and emerging invasive alien species in Europe. Global Change Biology 26:6235–6250.

Brook BW, Sodhi NS, Bradshaw CJA. 2008. Synergies among extinction drivers under global change. Trends in Ecology & Evolution 23:453–460.

Briski E et al. 2024. Does non-native diversity mirror Earth’s biodiversity? Global Ecology and Biogeography 33:48–62.

Carboni M, Calderon-Sanou I, Pollock L, Violle C, DivGrass Consortium, Thuiller W. 2018. Functional traits modulate the response of alien plants along abiotic and biotic gradients. Global Ecology and Biogeography 27:1173–1185.

Carrasco LR, Mumford JD, MacLeod A, Harwood T, Grabenweger G, Leach AW, Knight JD, Baker RHA. 2010. Unveiling human-assisted dispersal mechanisms in invasive alien insects: Integration of spatial stochastic simulation and phenology models. Ecological Modelling 221:2068–2075.

Carter ZT, Lumley T, Bodey TW, Russell JC. 2021. The clock is ticking: Temporally prioritizing eradications on islands. Global Change Biology 27:1443–1456.

Chalkowski K, Lepczyk CA, Zohdy S. 2018. Parasite Ecology of Invasive Species: Conceptual Framework and New Hypotheses. Trends in Parasitology 34:655–663.

Christie AP, Amano T, Martin PA, Petrovan SO, Shackelford GE, Simmons BI, Smith RK, Williams DR, Wordley CFR, Sutherland WJ. 2021. The challenge of biased evidence in conservation. Conservation Biology 35:249–262.

Clout MN, Russell JC. 2008. The invasion ecology of mammals: a global perspective. Wildlife Research 35:180–184.

Crowl TA, Crist TO, Parmenter RR, Belovsky G, Lugo AE. 2008. The spread of invasive species and infectious disease as drivers of ecosystem change. Frontiers in Ecology and the Environment 6:238–246.

Cuthbert R, Bodey TW, Briski E, Capellini I, Dick JTA, Kourantidou M, Ricciardi A, Pincheira-Donoso D. 2023, April 26. Harnessing Trait Evolution to Predict Economic Costs of Biological Invasions. Rochester, NY. Available from https://papers.ssrn.com/abstract=4430070

Cuthbert RN et al. 2022. Biological invasion costs reveal insufficient proactive management worldwide. Science of The Total Environment 819:153404.

Cuthbert RN, Dick JTA, Haubrock PJ, Pincheira-Donoso D, Soto I, Briski E. 2024. Economic impact disharmony in global biological invasions. Science of The Total Environment 913:169622.

Dana ED, Jeschke JM, García-de-Lomas J. 2014. Decision tools for managing biological invasions: existing biases and future needs. Oryx 48:56–63.

Diagne C, Ballesteros-Mejia L, Cuthbert RN, Bodey TW, Fantle-Lepczyk J, Angulo E, Bang A, Dobigny G, Courchamp F. 2023. Economic costs of invasive rodents worldwide: the tip of the iceberg. PeerJ 11:e14935.

Diagne C, Leroy B, Gozlan RE, Vaissière A-C, Assailly C, Nuninger L, Roiz D, Jourdain F, Jarić I, Courchamp F. 2020. InvaCost, a public database of the economic costs of biological invasions worldwide. Scientific Data 7:277.

Diagne C, Leroy B, Vaissière A-C, Gozlan RE, Roiz D, Jarić I, Salles J-M, Bradshaw CJA, Courchamp F. 2021. High and rising economic costs of biological invasions worldwide. Nature 592:571–576.

Dick JTA et al. 2017. Invader Relative Impact Potential: a new metric to understand and predict the ecological impacts of existing, emerging and future invasive alien species. Journal of Applied Ecology 54:1259–1267.

Doherty TS, Glen AS, Nimmo DG, Ritchie EG, Dickman CR. 2016. Invasive predators and global biodiversity loss. Proceedings of the National Academy of Sciences 113:11261– 11265.

Drake DR, Bodey TW, Russell JC, Towns DR, Nogales M, Ruffino L. 2011. Direct impacts of seabird predators on island biota other than seabirds. Pages 91-132 in Mulder CPH, Anderson WB, Towns DR, Bellingham PF editors. Seabird islands: Ecology, invasion and restoration. Oxford University Press, Oxford

Duncan RP. 2021. Time lags and the invasion debt in plant naturalisations. Ecology Letters 24:1363–1374.

Early R et al. 2016. Global threats from invasive alien species in the twenty-first century and national response capacities. Nature Communications 7:12485.

Essl F et al. 2020. Drivers of future alien species impacts: An expert-based assessment. Global Change Biology 26:4880–4893.

Evans T, Angulo E, Diagne C, Kumschick S, Şekercioğlu ÇH, Turbelin A, Courchamp F. 2023. Identifying links between the biodiversity impacts and monetary costs of alien birds. People and Nature 5:1561–1576.

Evans T, Blackburn TM, Jeschke JM, Probert AF, Bacher S. 2020. Application of the Socio-Economic Impact Classification for Alien Taxa (SEICAT) to a global assessment of alien bird impacts. NeoBiota 62:123–142.

Faurby S, Davis M, Pedersen RØ, Schowanek SD, Antonelli1 A, Svenning J-C. 2018. PHYLACINE 1.2: The Phylogenetic Atlas of Mammal Macroecology. Ecology 99:2626–2626.

Fournier A, Penone C, Pennino MG, Courchamp F. 2019. Predicting future invaders and future invasions. Proceedings of the National Academy of Sciences 116:7905–7910.

Gippet JMW, Bertelsmeier C. 2021. Invasiveness is linked to greater commercial success in the global pet trade. Proceedings of the National Academy of Sciences 118:e2016337118.

Hanley N, Roberts M. 2019. The economic benefits of invasive species management. People and Nature 1:124–137.

Henry M et al. 2023. Unveiling the hidden economic toll of biological invasions in the European Union. Environmental Sciences Europe 35:43.

Heringer G, Angulo E, Ballesteros-Mejia L, Capinha C, Courchamp F, Diagne C, Duboscq-Carra VG, Nuñez MA, Zenni RD. 2021. The economic costs of biological invasions in Central and South America: a first regional assessment. NeoBiota 67:401–426.

Hulme PE et al. 2008. Grasping at the routes of biological invasions: a framework for integrating pathways into policy. Journal of Applied Ecology 45:403–414.

Jeschke JM, Strayer DL. 2006. Determinants of vertebrate invasion success in Europe and North America. Global Change Biology 12:1608–1619.

Jetz W, Pyron RA. 2018. The interplay of past diversification and evolutionary isolation with present imperilment across the amphibian tree of life. Nature Ecology & Evolution 2:850–858.

Jetz W, Thomas GH, Joy JB, Hartmann K, Mooers AO. 2012. The global diversity of birds in space and time. Nature 491:444–448.

Jones HP et al. 2016. Invasive mammal eradication on islands results in substantial conservation gains. Proceedings of the National Academy of Sciences 113:4033–4038.

Jones BA. 2017. Invasive Species Impacts on Human Well-being Using the Life Satisfaction Index. Ecological Economics 134:250–257.

Kaiser BA, Burnett KM. 2006. Economic impacts of E. Coqui frogs in Hawaii. Interdisciplinary Environmental Review 8:1–11.

King CM. 2017. Pandora’s box down-under: origins and numbers of mustelids transported to New Zealand for biological control of rabbits. Biological Invasions 19:1811–1823.

Kourantidou M, Haubrock PJ, Cuthbert RN, Bodey TW, Lenzner B, Gozlan RE, Nuñez MA, Salles J-M, Diagne C, Courchamp F. 2022. Invasive alien species as simultaneous benefits and burdens: trends, stakeholder perceptions and management. Biological Invasions 24:1905–1926.

Kraus F. 2015. Impacts from Invasive Reptiles and Amphibians. Annual Review of Ecology, Evolution, and Systematics 46:75–97.

Kumschick S et al. 2015. Ecological Impacts of Alien Species: Quantification, Scope, Caveats, and Recommendations. BioScience 65:55–63.

Langhammer PF et al. 2024. The positive impact of conservation action. Science 384:453–458.

Latombe G, Catford JA, Essl F, Lenzner B, Richardson DM, Wilson JRU, McGeoch MA. 2022. GIRAE: a generalised approach for linking the total impact of invasion to species’ range, abundance and per-unit effects. Biological Invasions 24:3147–3167.

Laughlin DC. 2014. Applying trait-based models to achieve functional targets for theory-driven ecological restoration. Ecology Letters 17:771–784.

Leroy B, Kramer AM, Vaissière A-C, Kourantidou M, Courchamp F, Diagne C. 2022. Analysing economic costs of invasive alien species with the invacost r package. Methods in Ecology and Evolution 13:1930–1937.

Leung B et al. 2012. TEASIng apart alien species risk assessments: a framework for best practices. Ecology Letters 15:1475–1493.

Lever C. 1985. Naturalized mammals of the world. Naturalized mammals of the world. Longman.

Lodge DM et al. 2016. Risk Analysis and Bioeconomics of Invasive Species to Inform Policy and Management. Annual Review of Environment and Resources 41:453–488.

Marino C, Bellard C. 2023. When origin, reproduction ability and diet define the role of birds in invasions. Proceedings of the Royal Society B: Biological Sciences 290:20230196.

Marino C, Leclerc C, Bellard C. 2022. Profiling insular vertebrates prone to biological invasions: What makes them vulnerable? Global Change Biology 28:1077–1090.

McCreless EE, Huff DD, Croll DA, Tershy BR, Spatz DR, Holmes ND, Butchart SHM, Wilcox C. 2016. Past and estimated future impact of invasive alien mammals on insular threatened vertebrate populations. Nature Communications 7:12488.

Meerburg BG, Singleton GR, Kijlstra A. 2009. Rodent-borne diseases and their risks for public health. Critical Reviews in Microbiology 35:221–270.

Myhrvold NP, Baldridge E, Chan B, Sivam D, Freeman DL, Ernest SKM. 2015. An amniote life-history database to perform comparative analyses with birds, mammals, and reptiles. Ecology 96:3109–3109.

Nuñez MA, Chiuffo MC, Pauchard A, Zenni RD. 2021. Making ecology really global. Trends in Ecology & Evolution 36:766–769.

Oppel S, Beaven BM, Bolton M, Vickery J, Bodey TW. 2011. Eradication of Invasive Mammals on Islands Inhabited by Humans and Domestic Animals. Conservation Biology 25:232–240.

Pili AN et al. 2024. Forecasting potential invaders to prevent future biological invasions worldwide. Global Change Biology 30:e17399.

Pigot AL, Sheard C, Miller ET, Bregman TP, Freeman BG, Roll U, Seddon N, Trisos CH, Weeks BC, Tobias JA. 2020. Macroevolutionary convergence connects morphological form to ecological function in birds. Nature Ecology & Evolution 4:230–239.

Pincheira-Donoso, D. 2017. The Global Amphibian Biodiversity Project (GABiP): an online facility for scientific information on the biodiversity, ecology and evolution of amphibians. https://www.amphibianbiodiversity.org.

Polaina E, Pärt T, Recio MR. 2020. Identifying hotspots of invasive alien terrestrial vertebrates in Europe to assist transboundary prevention and control. Scientific Reports 10:11655.

Pyšek P et al. 2020. Scientists’ warning on invasive alien species. Biological Reviews 95:1511–1534.

Pyšek P, Richardson DM. 2010. Invasive Species, Environmental Change and Management, and Health. Annual Review of Environment and Resources 35:25–55.

Ricciardi A et al. 2021. Four priority areas to advance invasion science in the face of rapid environmental change. Environmental Reviews 29:119–141.

Ricciardi A, Hoopes MF, Marchetti MP, Lockwood JL. 2013. Progress toward understanding the ecological impacts of nonnative species. Ecological Monographs 83:263–282.

Rodda GH, Savidge JA. 2007. Biology and Impacts of Pacific Island Invasive Species. 2. Boiga irregularis, the Brown Tree Snake (Reptilia: Colubridae)1. Pacific Science 61:307–324.

Roy HE et al. 2024. IPBES Invasive Alien Species Assessment: Summary for Policymakers. Zenodo. Available from https://zenodo.org/records/10521002

Russell JC, Broome KG. 2016. Fifty years of rodent eradications in New Zealand: another decade of advances. New Zealand Journal of Ecology 40:197–204.

Scheele BC et al. 2019. Amphibian fungal panzootic causes catastrophic and ongoing loss of biodiversity. Science 363:1459–1463.

Seebens H et al. 2017. No saturation in the accumulation of alien species worldwide. Nature Communications 8:14435.

Seebens H et al. 2020. A workflow for standardising and integrating alien species distribution data. NeoBiota 59:39–59.

Seebens H et al. 2021. Projecting the continental accumulation of alien species through to 2050. Global Change Biology 27:970–982.

Shackleton RT et al. 2019. Explaining people’s perceptions of invasive alien species: A conceptual framework. Journal of Environmental Management 229:10–26.

Sol D, Bacher S, Reader SM, Lefebvre L. 2008. Brain Size Predicts the Success of Mammal Species Introduced into Novel Environments. The American Naturalist 172:S63–S71.

Sol D, Maspons J, Vall-llosera M, Bartomeus I, García-Peña GE, Piñol J, Freckleton RP. 2012. Unraveling the Life History of Successful Invaders. Science 337:580–583.

Soto I et al. 2024a. The wild cost of invasive feral animals worldwide. Science of The Total Environment 912:169281.

Soto I et al. 2024b. Taming the terminological tempest in invasion science. Biological Reviews 99:1357–1390.

Soto I, Cuthbert RN, Kouba A, Capinha C, Turbelin A, Hudgins EJ, Diagne C, Courchamp F, Haubrock PJ. 2022. Global economic costs of herpetofauna invasions. Scientific Reports 12:10829.

Spatz DR, Zilliacus KM, Holmes ND, Butchart SHM, Genovesi P, Ceballos G, Tershy BR, Croll DA. 2017. Globally threatened vertebrates on islands with invasive species. Science Advances 3:e1603080.

Steffen W et al. 2015. Planetary boundaries: Guiding human development on a changing planet. Science 347:1259855.

Stobo-Wilson AM et al. 2022. Counting the bodies: Estimating the numbers and spatial variation of Australian reptiles, birds and mammals killed by two invasive mesopredators. Diversity and Distributions 28:976–991.

Stone GN, Nee S, Felsenstein J. 2011. Controlling for non-independence in comparative analysis of patterns across populations within species. Philosophical Transactions of the Royal Society B: Biological Sciences 366:1410–1424.

Street SE, Gutiérrez JS, Allen WL, Capellini I. 2023. Human activities favour prolific life histories in both traded and introduced vertebrates. Nature Communications 14:262.

Streit RP, Bellwood DR. 2023. To harness traits for ecology, let’s abandon ‘functionality.’ Trends in Ecology & Evolution 38:402–411.

Tobias JA et al. 2022. AVONET: morphological, ecological and geographical data for all birds. Ecology Letters 25:581–597.

Tonini JFR, Beard KH, Ferreira RB, Jetz W, Pyron RA. 2016. Fully-sampled phylogenies of squamates reveal evolutionary patterns in threat status. Biological Conservation 204:23–31.

Toomes A, García-Díaz P, Stringham OC, Ross JV, Mitchell L, Cassey P. 2022. Drivers of the Australian native pet trade: The role of species traits, socioeconomic attributes and regulatory systems. Journal of Applied Ecology 59:1268–1278.

Turbelin AJ et al. 2022. Introduction pathways of economically costly invasive alien species. Biological Invasions 24:2061–2079.

Uetz, P., Freed, P, Aguilar, R., Reyes, F., Kudera, J. & Hošek, J. 2023. The Reptile Database, http://www.reptile-database.org

Upham NS, Esselstyn JA, Jetz W. 2019. Inferring the mammal tree: Species-level sets of phylogenies for questions in ecology, evolution, and conservation. PLOS Biology 17:e3000494.

Vaissière A-C, Courtois P, Courchamp F, Kourantidou M, Diagne C, Essl F, Kirichenko N, Welsh M, Salles J-M. 2022. The nature of economic costs of biological invasions. Biological Invasions 24:2081–2101.

Veitch CR, Clout MN, Martin AR, Russell JC, West CJ, editors. 2019. Island invasives: scaling up to meet the challenge. Proceedings of the international conference on island invasives 2017. IUCN, International Union for Conservation of Nature.

Wang S, Deng T, Zhang J, Li Y. 2023. Global economic costs of mammal invasions. Science of The Total Environment 857:159479.

Wang S, Li W, Zhang J, Luo Z, Li Y. 2024. Alien range size, habitat breadth, origin location, and domestication of alien species matter to their impact risks. Integrative Zoology.

Wilman H, Belmaker J, Simpson J, de la Rosa C, Rivadeneira MM, Jetz W. 2014. EltonTraits 1.0: Species-level foraging attributes of the world’s birds and mammals. Ecology 95:2027– 2027.

Witmer GW, Shiels AB. 2017. Ecology, Impacts, and Management of Invasive Rodents in the United States. Pages 193–220 in Pitt WC, editor. Ecology and Management of Terrestrial Vertebrate Invasive Species in the United States, 1st edition. CRC Press, Boca Raton: Taylor & Francis, 2018.

Wood SN. 2011. Fast stable restricted maximum likelihood and marginal likelihood estimation of semiparametric generalized linear models. Journal of the Royal Statistical Society: Series B 73:3–36.

